# After migration into blood circulation, hematopoietic stem cells can stably self-renew and maintain bone marrow while being skewed toward myeloid lineage when submitted to serial transplantation

**DOI:** 10.1101/2021.03.04.433880

**Authors:** Yasmine Even, Lin Yi, Chih-Kai Chang, Fabio M. Rossi

**Affiliations:** Department of Medicine, The Biomedical Research Centre, University of British Columbia, Vancouver, BC, V6T1Z3, Canada; LEMAR, Institut Universitaire Européen de la Mer, Université de Bretagne Occidentale, Plouzané, 29280, France

**Keywords:** hematopoietic stem cells, migration, self-renewal, transplantation

## Abstract

Hematopoietic stem cells (HSCs) mainly reside in bone marrow (BM) within niches providing an appropriate environment for their survival and self-renewal. Although, small numbers of HSCs can quit their residing environment to migrate into blood circulation and re-engraft elsewhere in BM. Mobilizing agents such as granulocyte colony stimulating factor (G-CSF) can amplify this process by inducing massive HSC mobilization into blood circulation. This method is widely used in clinics to treat hematological disorders. However, in physiological conditions, the properties of HSCs after migration (called migratory HSCs) remain incompletely characterized. In this study, we investigated the capacity of migratory HSCs to self-renew, reconstitute and maintain BM. We show that after migration, HSCs can stably self-renew and maintain BM in homeostasis. However, while stably repopulating BM of irradiated recipients, migratory HSCs show a defect in lymphoid lineage reconstitution when subjected to serial transplantations. Our findings provide interesting knowledge on HSC properties after migration, which may benefits therapeutic research on HSC-based therapies to treat hematological disorders.

## Introduction

Hematopoietic stem cells (HSCs) mainly reside in bone marrow (BM) within HSC niches where the microenvironment supports their maintenance and self-renewal [1]. It is well known that in certain conditions, such as after a G-CSF treatment, a massive number of HSCs can migrate into blood circulation [2,3]. This process of HSC mobilization is widely used in clinics to provide hematopoietic grafts [4,5]. In addition, several studies have demonstrated that HSCs can leave their residential environment to migrate into blood circulation during homeostasis [6,7]. In a more recent work, about 1-5% of HSCs have been estimated to enter the bloodstream every day in healthy mice [8]. Despite the fact that this phenomenon has been first described over 50 years ago, its regulation and purpose are still poorly understood [9,10,11]. It has been hypothesized that HSC circulation may be important for peripheral immune surveillance [12]. Based on parabiosis experiments, another group has shown that circulating HSCs can stably relocate to the BM and suggested that migration would serve to maintain even HSC numbers across this diffused compartment [6]. However other parabiosis-based studies suggest that after migration across the parabiotic junction into the mouse partner, HSCs do not stably return to BM, have impaired homing ability and only transiently repopulate BM, casting doubts on whether they can efficiently self-renew [7]. This raises the question whether migration would change intrinsic HSC properties. A better knowledge on HSCs after migration into bloodstream would be useful to improve clinical uses of HSCs for hematological disorder treatments. Therefore, in the current study, we used parabiosis to investigate HSC characteristics before and after migration in physiological conditions as well as during myelotoxic stress.

## Materials and methods

### Animals

All mice were bred in-house and maintained in a pathogen-free environment. Heterozygous GFP^+^ Ly5.2 C57BL/6 females, expressing GFP ubiquitously from the cytomegalovirus enhancer-chicken β-actin hybrid promoter [13], and their non-GFP littermate sisters were used for parabiosis. C57BL/6 Ly5.1 female mice, 8-12 weeks of age, were used as recipients. All experiments were approved by the University of British Columbia Animal Care Committee (Protocol Numbers: A05-0351 and A06-0185).

### Antibodies for flow cytometry analyses

Anti-CD3 conjugated to PE was purchased from BD Biosciences Pharmingen (San Diego, CA); anti-B220 conjugated to PE, anti-Gr-1 and anti-Mac1 conjugated to PE-Cy7 and anti-CD45.2 conjugated to APC were purchased from eBioscience (San Diego, CA).

### Parabiosis and separation surgery

Pairs of 8-to 12-week-old female littermates were housed together in a single cage for 2 weeks, and then joined in parabiosis. Mice were anesthetized with isoflurane (1.5-2%, inhaled) and joined as described in [2,6]. Briefly, on one side of each mouse, skin was incised, olecranon and knee joints were attached by suture and tie and mouse skins were sutured by staples. Parabionts were surgically separated 6 weeks after parabiosis surgery by a reversal of the above procedure.

### Bone marrow harvest for analysis and transplantation

BM was harvested using the identical method for each experiment: whole BM from mice was obtained by flushing femur and tibia.

### Flow cytometry analysis of migratory HSCs in separated parabionts

After a period of 4 and 6 months post-separation, the proportion of BM HSCs coming from each partner was quantified by flow cytometry (FACSCalibur, BD Biosciences). To do so, blood and BM from separated parabionts were harvested and single cell suspensions were stained for Gr1 to select the short lived granulocyte population and CD3 to exclude any long-lived lymphoid cell. Granulocyte having a short half-life (∼3 days), after 4 months of separation all granulocytes that migrated into the partner mouse or that have issued from short-term HSCs and progenitor cells have disappeared. Therefore any GFP^+^ granulocyte (Gr1^+^CD3^-^) found in BM or peripheral blood of a non-GFP mouse must have issued from long-term HSCs that have migrated and re-engrafted the partner’s BM (and vice versa: GFP^-^ granulocytes for a GFP mouse). Therefore, the proportion of GFP^+^ granulocytes should reflect the proportion of cross engrafted LT-HSCs in non-GFP separated parabiont and vice versa. To quantify differentiated lineage cells issued from HSCs, BM from separated parabionts was stained for Gr1, Mac1, CD3 and B220 and analyzed by flow cytometry (FACSCalibur, BD Biosciences).

### BM transplantations and flow cytometry analysis

Lethally irradiated (9.5 Gy) 8-to 12-week-old mice (Ly5.1) were used as primary and secondary recipients. Primary recipients received 5×10^6^ total BM cells from parabionts separated for 4 months. BM from each separated parabiont was injected into at least 3 recipients. Four, 6 and 12 months after transplantation, blood and BM from irradiated recipients were stained for Gr1, Mac1, CD3, B220 and CD45.2 markers and analyzed by flow cytometry (FACSCalibur, BD Biosciences) in order to quantify the participation of partner-derived (GFP^+^) HSCs in BM repopulation. All recipient mice exhibited >90% chimerism.

### Serial transplantation of migratory HSCs

Four months post-separation, 5×10^6^ total BM cells from parabionts exhibiting 2±0.3% cross-engrafted HSCs (GFP^+^) were intravenously injected into 3 irradiated primary Ly5.1 recipients. Primary recipients exhibiting 5±0.4% GFP^+^ BM cells were chosen as donors. Secondary recipients received 5×10^6^ total BM cells from selected primary recipients. Analysis by flow cytometry was performed 4 months post-transplantation as described above. Recipient BM and blood were harvested 4 months post-transplant and stained for myeloid (Gr1, Mac1) and lymphoid population (CD3, B220) and CD45.2. The relative contribution of donor cells issued from migratory HSCs to BM reconstitution was assessed by quantifying the proportion of GFP^+^ cells among the CD45.2^+^ lineage population.

### 2-98% BM competition assay (serial transplantation of resident HSCs)

Lethally irradiated (9.5 Gy) 8-to 12-week-old mice (Ly5.1) were intravenously injected with a total of 5×10^6^ total BM cells from 8-week-old donor mice. The transplanted cells comprised a 98:2 ratio of respectively, Ly5.2 and Ly5.2-GFP^+^ BM harvested from female littermates. Lethally irradiated secondary recipients were transplanted with 5×10^6^ total BM cells from primary recipients exhibiting 5±0.5% GFP^+^ BM cells and flow cytometry analysis was performed 4 months post-transplantation as described above. Only recipient mice exhibiting >90% chimerism (Ly5.2) were analyzed.

### Statistical methods

Data are presented as mean ± SEM. Data were analyzed using the Kruskal-Wallis test and differences were considered statistically significant for a P≤ 0.05.

## Results

### Following re-engraftment, migratory HSCs are stably maintained in BM

It is now well established that, in parabiotic animals, a small proportion of HSCs migrates through the circulation and cross-engrafts the partner’s BM [2,6,7]. However, the stability of such cross-engraftment in BM remains unclear. To directly address this issue, we joined in parabiosis two Ly5.2 littermates, one of which constitutively expressing GFP in all cells. We allowed migration to take place for 6 weeks, following which we surgically separated the parabionts. The proportion of cross-engrafted HSCs (coming from the partner and also called migratory HSCs) was quantified by flow cytometry 4 and 6 months after separation, based on GFP expression within the short-lived CD3^-^ Gr1^+^ granulocyte population (Figure 1A&B). Granulocytes are short term living cells and consequently usually chosen to measure the proportion of LT-HSCs in parabiosis. We found 3% partner-derived granulocytes issued from cross-engrafted HSCs in parabionts 4 months after separation (Figure 1 B&C). This result, reflecting the presence of 3% HSCs engrafted in the BM after migration is in accordance with previous studies. An additional 2 months later the proportion of cross-engrafted HSCs did not change, suggesting that migrating HSCs can stably re-engraft BM over a period of at least 6 months (Figure 1C). Furthermore, in order to make sure that LT-HSCs contributed to all blood cell lineages, we also quantified the proportion of myeloid, B and T cells issued from migratory HSCs. We call them partner-derived cells because blood cell lineages might include long-lived cells, especially memory T cells. As shown on figure 1D lymphoid and myeloid populations mainly issued from the migratory HSCs are similar at 4 and 6 months post-separation indicating that migratory HSCs contributed to all differentiated blood cell lineages at 4 and 6 months post separation.

**Figure 1.**
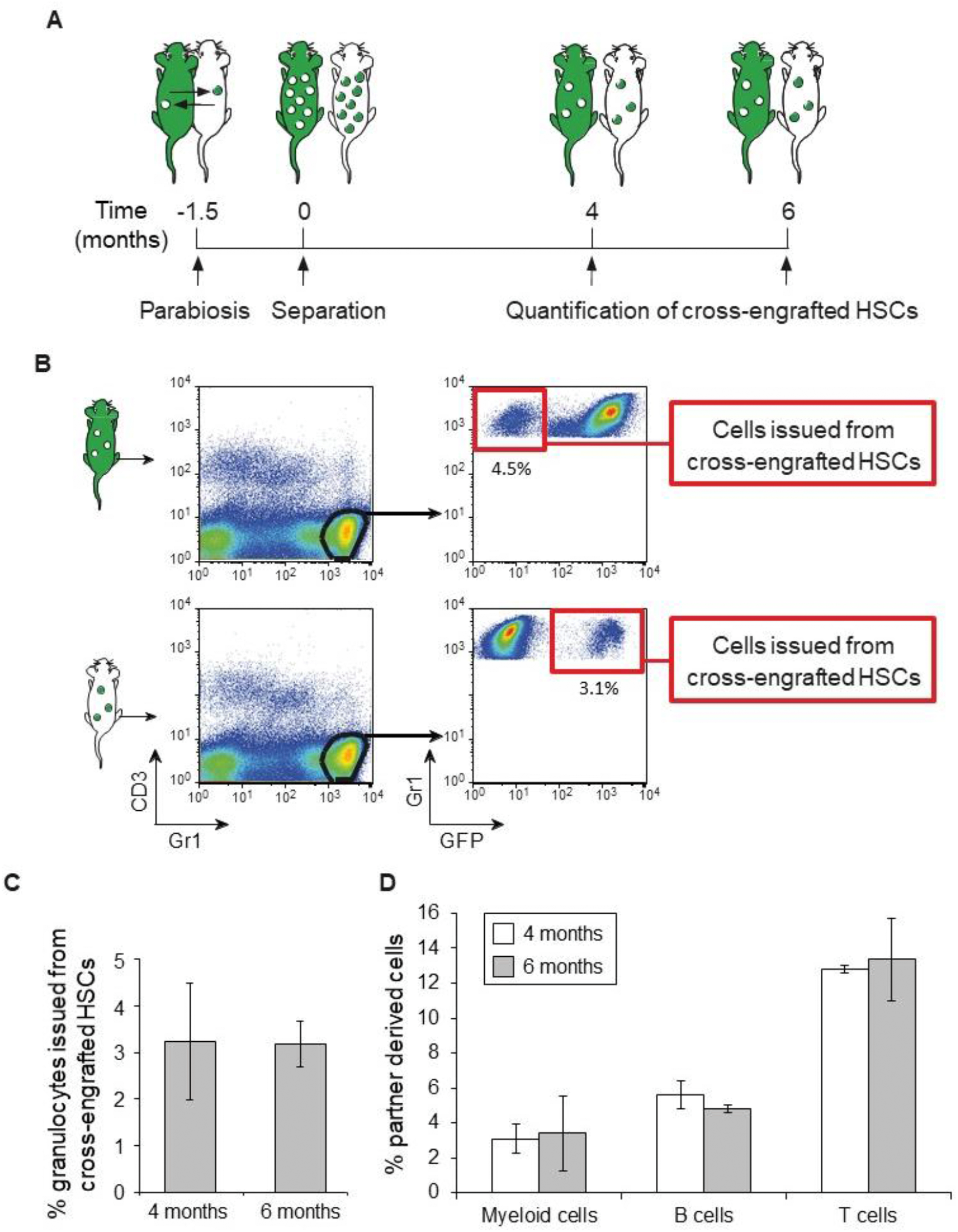
HSCs can migrate into blood circulation and stably re-engraft BM. A. Schematic of the experimental design. Parabiosis leads to peripheral blood exchange, allowing cells to migrate from one partner to the other one. B, C. BM of separated parabionts was harvested and the frequency of granulocytes derived from cross-engrafted HSCs was measured by flow cytometry. B. Dot plots show representative analysis of a GFP (upper plots) and a non-GFP partner (lower plots). Gated regions designate granulocytes from BM (left plots) and granulocytes issued from partner-derived (cross-engrafted, also called migratory) HSCs (right plots). C. Histogram shows the proportion of granulocytes issued from partner-derived (cross-engrafted) HSCs 4 and 6 months post-separation (N=3). D. Histogram shows the proportion of myeloid, B and T cells that are partner derived (which means cells that are issued from migratory HSCs + potential remaining long-lived cells (N=3)).

### After migration, HSCs keep self-renewal ability over a period of at least 6 months

In order to assess the self-renewal potential of HSCs after migration, we transplanted BM from each separated non-GFP parabiont into 3 lethally irradiated recipients (Figure 2A). Four, 6 and 12 months after transplantation, we measured the ability of cross-engrafted HSCs (also called migratory HSCs) to reconstitute all differentiated peripheral blood lineages (Figures 2B&C). We found that the proportion of myeloid, B- and T-cell populations issued from migratory HSCs in primary recipients averaged 2% over the course of our analysis (figure 2C). However, a higher variance can be observed at 12 months for all 3 differentiated lineages. More precisely, the proportion of each differentiated lineage issued from migratory HSCs is over 0.5% in almost all recipients’ BM at 4 and 6 months but only in half of recipients’ BM at 12 months. This result suggests that at 12 months post-transplantation, part of the pool of migratory HSCs that have engrafted the BM may not fulfill the function of BM maintenance any more, possibly due to exhaustion, decreased proliferation or differentiation into more mature progenitors.

**Figure 2.**
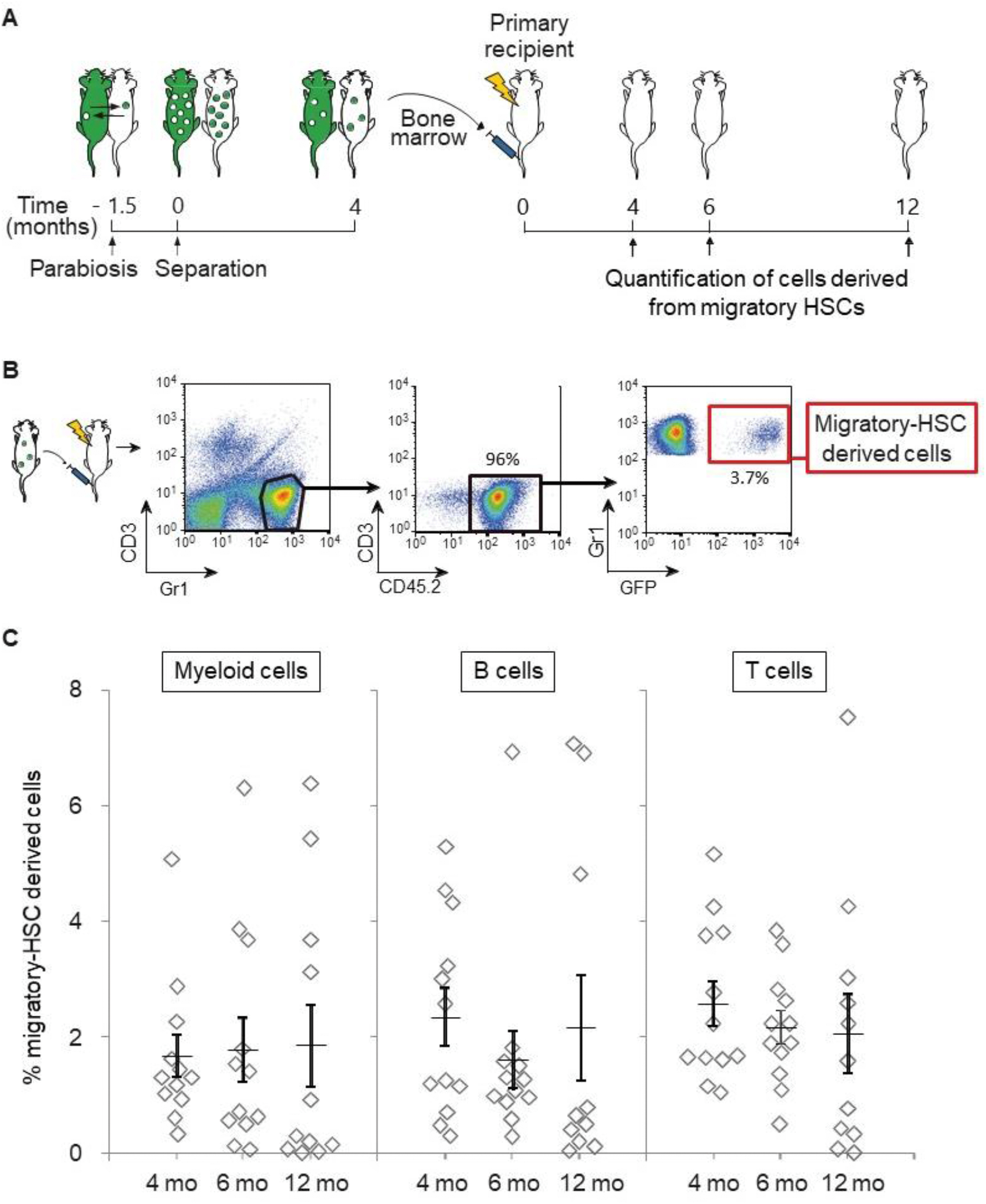
Cross-engrafted HSCs can repopulate and maintain BM of irradiated recipient for at least 6 months. A. Schematic of the experimental design. B, C. The peripheral blood and BM of recipient mice were harvested and among donor-derived cells (CD45.2^+^), the frequency of myeloid cells (Gr1^+^ and Mac1^+^), T cells (CD3^+^) and B cells (B220^+^) derived from cross-engrafted HSCs (GFP^+^) was measured by flow cytometry. B. Dot plots show representative analyses of recipient BM. Gated regions designate granulocytes (left plot), donor-derived granulocytes (middle plot) and granulocytes issued from partner-derived (migratory) HSCs (right plot). C. Graphs show the frequency of differentiated cells issued from partner-derived (migratory) HSCs, at 4, 6 and 12 months post-transplantation. Each diamond represents a recipient mouse. Each horizontal line indicates the mean percent (± SEM) of at least 10 recipients for each lineage and time point.

### Differentiation of migratory HSCs is skewed toward myeloid lineage when submitted to serial transplantation

In order to better define BM reconstitution properties of migratory HSCs, we compared them to resident HSCs in equivalent experimental conditions within 2 rounds of serial transplantation. To do so, irradiated primary recipients (Ly 5.1) were transplanted either with separated-parabiont BM containing 2% cross-engrafted HSCs (GFP^+^) (Figure 3A left part, migratory-HSC group) or with a mixture of 2% Ly5.2-GFP BM and 98% Ly5.2 BM (Figure 3A left part, resident-HSC group). Thereafter, primary recipients (resident- or migratory-HSC injected recipients) exhibiting 5±0.5% GFP^+^-HSCs where transplanted into irradiated secondary recipients (Figure 3A right part). We observed that the average of myeloid cells stemming from GFP^+^ resident BM HSCs was not significantly different from the one of migratory BM HSCs in primary and secondary recipients (Figures 3B&C). Furthermore, the proportion of lymphoid cells issued from migratory and resident HSCs was the same in primary recipients (Figure 3B). However, in secondary recipients, there was a significantly lower rate of lymphoid cells issued from GFP^+^ HSCs in the migratory-HSC group than in the resident-HSC group (Figure 3C). Overall, these results suggest that after leaving the BM niche, migrating through the blood circulation and re-engrafting the BM, HSCs may be skewed toward the myeloid lineage when subjected to increased self-renewal divisions.

**Figure 3.**
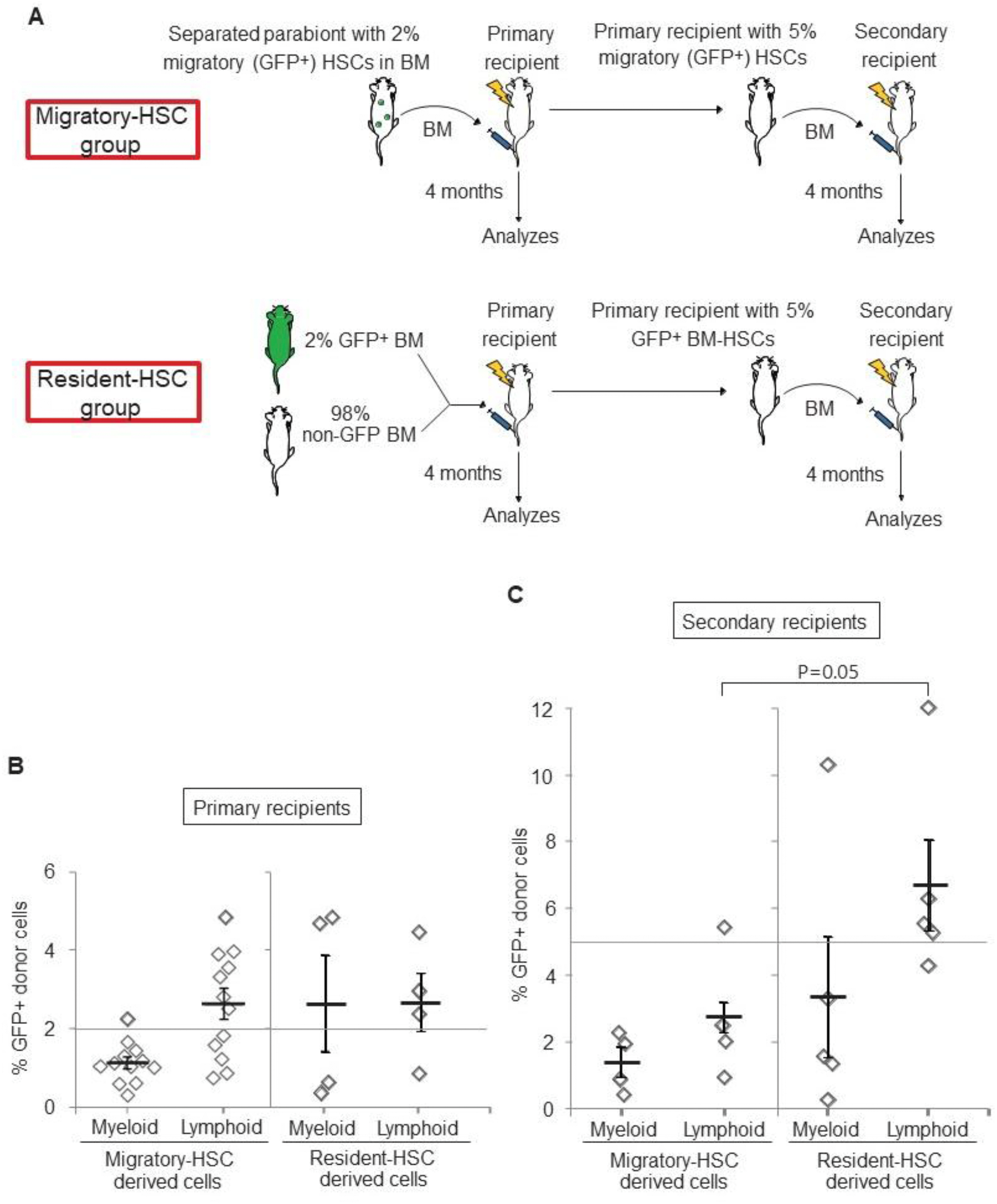
Myeloid lineage skewing of migratory HSCs in high cell division rate conditions. A. Schematic of the experimental design: migratory-HSC group (left part). Separated parabionts exhibiting 2±0.3% migratory GFP^+^-HSCs in BM were used as donor mice for irradiated primary recipients. Resident-HSC group (left part). Ratios of 98% non-GFP and 2% GFP^+^ BM cells were injected into irradiated primary recipients. A (right part). BM of primary recipients reconstituted with a proportion of 5±0.5% migratory or resident BM GFP^+^-HSCs were injected into irradiated secondary recipients. B&C. Graphs show the proportion of myeloid and lymphoid cells issued from migratory or resident GFP^+^-HSCs in primary and secondary recipients 4 months post-transplantation. Each diamond represents a mouse and each horizontal line indicates the mean percent (± SEM) of at least 4 mice. Dashed line shows the proportion of GFP^+^-HSCs in donor BM.

## Discussion

In our study we show that HSCs can migrate through blood circulation, re-engraft BM and generate progeny for at least 6 months in homeostatic conditions. In addition, migratory HSCs can reconstitute and stably maintain the BM of irradiated recipients for at least 6 months while starting to lose repopulation ability at one year post-transplantation. We further demonstrate that after migration, HSCs are less competent in reconstituting the lymphoid population than resident HSCs in serially transplanted animals.

We first show that 3% cross-engrafted HSCs are present in parabiont BM, 4 and 6 months after separation comforting the results of previous parabiosis-based studies demonstrating stable re-engraftment [6]. The low numbers of cross-engrafted HSCs is also supported by a recent study showing that some HSCs exhibit limited migration toward other niches [14]. It is to note that there may be a fraction of migratory HSCs that are quiescent and not contributing to hematopoiesis. Therefore, our data that are based on HSCs participating in hematopoiesis, may underestimate the proportion of migratory HSCs. Moreover, the parabiosis-based model involves surgery. Therefore, we also have to consider that the transient release of inflammatory cytokines and chemokines induced by the intervention may increase HSC egress from the BM to blood circulation. Therefore our results may not completely reflect what is happening in physiological conditions. We further showed that cross-engrafted migratory-HSCs stably self-renewed and repopulated recipient BM until at least 6 months post-transplantation, illustrating that HSCs keep their self-renewal ability after migration in accordance with previous studies [6]. However, these results are in contrast to studies suggesting that migrating HSCs only transiently re-engraft the BM in both physiological conditions and irradiated recipients [7]. In comparison with this study, we injected 20 times more parabionts’ BM cells (5×10^6^ versus 2.5×10^5^) into irradiated recipients, which might partly explain the differences in the results. Indeed, considering the low percentage of migratory HSCs found in partner BM, so few injected BM cells likely led to limiting numbers of stem cells being transplanted [15].

We also observed that the proportion of migratory HSCs capable of BM maintenance was decreased at one year post-transplantation with higher standard error suggesting higher variability of migratory-HSC property to reconstitute BM. We further demonstrate that, while keeping their long-term repopulation ability, migratory HSCs are less competent in reconstituting the lymphoid population than resident HSCs in serially transplanted animals. Based on single cell transplantations, several studies have shown variable self-renewal properties within the HSC population [14,16,17]. Indeed McKenzie et al. found that the human HSC population comprises stable clones capable of self-renewal until at least 7 months as well as other clones that varied in their contribution to hematopoiesis over time. Dykstra et al. also found distinct clones with more specific role among the HSC population. More precisely they identified 4 distinct BM HSC subpopulations with a myeloid or lymphoid bias that could be observed in secondary and tertiary recipients. This raises the question whether migratory HSCs showing myeloid bias would constitute one of these identified populations. Furthermore, other studies have demonstrated that depending on their location in the bone marrow HSCs do not behave equally in term of regenerative and self-renewal capacities [18,19]. Therefore, it would be interesting to know whether migratory HSCs come from a specific location in BM. Furthermore, since the difference between lymphoid and myeloid lineage from resident and migratory HSCs could be the result of a differentiation defect or of cell exhaustion, further analyses on progenitors issued from migratory HSCs could give more insight into potential mechanism of this phenomenon.

Our findings that migratory and resident HSCs behave differently could also be explained by possible different function to play. Whereas it is essential for resident HSCs to maintain BM integrity by self-renewing and producing all types of blood cells. Migratory HSCs would further serve a different role such as immune survey or tissue repair. Indeed, Massberg et al. have described that HSCs and early blood progenitors can transit into the lymphatic system and then may return to BM via the blood circulation to produce differentiated blood cells. The authors suggest that HSCs migrate to provide a rapid respond unit in peripheral tissues involved in immune protection [12]. And in a more recent study, it was shown that infection was activating HSC mobilization into blood circulation [20]. Therefore, several studies comfort the idea that circulating HSCs could exert a role of survey or repair damaged tissues.

Furthermore, we cannot exclude that part of migratory HSCs may relocate somewhere else than in the BM. Indeed it is well known that HSCs can migrate and seek for other extramedullary hematopoietic organs such as spleen or liver where they keep their HSC properties or they may differentiate into more specialized progenitors [6,21,22]. Furthermore, HSCs were identified in muscle tissues where they may participate in the healing process after tissue injury [7,23]. Further investigations on migratory HSCs such as marker identification and localization would help in their characterization.

In summary, our findings that HSCs behave similarly before and after migration in physiological conditions are in accordance with the theory that at any time in the body a small part of HSCs circulate through the bloodstream and re-engraft somewhere else within the BM compartment to maintain homogeneity and integrity of the whole BM. However, in myeloablative conditions migratory HSCs behave differently and may acquire other properties, which would support the proposal that migratory HSCs participate in peripheral immune surveillance and in tissue repair after injury. These findings provide interesting knowledge that may be useful to improve clinical treatments regarding hematological diseases.

## Acknowledgements

We thank J.-H. Kang and A. Krüll for technical help, the Biomedical Research Centre Animal Facility and core staff, and A. Johnson of the University of British Columbia Multi-User Flow Cytometry/Fluorescence-activated Cell Sorting Facility for their expert technical assistance. This work was supported by a CIHR grant MOP 160678 to F.M. Rossi. F.M. Rossi holds a Canada Research Chair in Regenerative Medicine. Y. Even was supported by a postdoctoral fellowship from la Fondation pour la Recherche Medicale (FRM, France).

